# Compound specific stable isotope analysis of amino acid nitrogen reveals detrital support of microphytobenthos in the Dutch Wadden Sea benthic food web

**DOI:** 10.1101/2022.05.24.493073

**Authors:** Philip M. Riekenberg, Tjisse van der Heide, Sander J. Holthuijsen, Henk W. van der Veer, Marcel T.J. van der Meer

**Affiliations:** Department of Marine Microbiology & Biogeochemistry, NIOZ Royal Netherlands Institute for Sea Research, PO Box 59, 1790AB Den Burg Texel, The Netherlands; Department of Coastal Systems, NIOZ Royal Netherlands Institute for Sea Research, PO Box 59, 1790AB Den Burg Texel, The Netherlands; Conservation Ecology Group, Groningen Institute for Evolutionary Life Sciences, University of Groningen, 9700 CC, Groningen, The Netherlands

**Keywords:** trophic discrimination, intertidal, diatoms, microbial loop, Wadden Sea, benthic food web, permeable sands

## Abstract

The Wadden Sea is the world’s largest intertidal ecosystem and provides vital food resources for a large number of migratory bird and fish species during seasonal stopovers. Previous work using bulk stable isotope analysis of carbon found that microphytobenthos was the dominant resource use fueling the food web with particulate organic matter making up the remainder. However, this work was unable to account for the trophic structure of the food web or the considerable increase in δ^15^N values of bulk tissue throughout the benthic food web occurring in the Eastern regions of the Dutch Wadden Sea. Here, we combine compound specific and bulk analytical stable isotope techniques to further resolve the trophic structure and resource use throughout the benthic food web in the Wadden Sea. Analysis of δ^15^N for trophic and source amino acids allowed for better identification of trophic relationships due to the integration of underlying variation in the nitrogen resources supporting the food web. Baseline integrated trophic position estimates using glutamic acid (Glu) and phenylalanine (Phe) allow for disentanglement of baseline variations in underlying δ^15^N sources supporting the ecosystem and trophic shifts resulting from changes in ecological relationships. Through this application we further confirmed dominant ecosystem support by microphytobenthos derived resources, although to a lesser extent than previously estimated. In addition to phytoplankton derived particulate organic matter and microphytobenthos supported from nutrients from the overlying water column there appears to be an additional resource supporting the benthic community. From the stable isotope mixing models, a subset of species appears to focus on microphytobenthos supported off recycled (porewater) N and/or detrital organic matter mainly driven by increased phenylalanine δ^15^N values. This additional resource within microphytobenthos may play a role in subsidizing the exceptional benthic productivity observed within the Wadden Sea ecosystem and reflect division in microphytobenthos support along green (herbivory) and brown (recycled/detrital) food web pathways.

## Introduction

The Wadden Sea is the world’s largest intertidal ecosystem (Wolff 1983) stretching from The Netherlands to Denmark formed behind a barrier island chain with connection to the North Sea (Postma 1996). This intertidal ecosystem has been designated as a culturally significant UNESCO World Heritage site due to considerable biodiversity within the system as benthic productivity supports an estimated 10 – 12 million migratory birds across each year (Reise, et al. 2010). The Wadden Sea has a long history of multiple direct impacts from human activity (Wolff 2013, Eriksson, et al. 2010) that includes land reclamation, partial damming and hydraulic changes, eutrophication, overfishing, and extensive dredging for shellfish. These impacts have resulted in a shift from a benthos dominated by seagrass, extensive bivalve reefs, and supporting apex predators towards one dominated by polychaetes, with minimal fringing bivalve reefs, and largely devoid of apex predators (Philippart, et al. 2007). Despite historical and recent anthropogenic impacts, the Wadden Sea remains a very productive intertidal ecosystem. However, changes in higher trophic level species have coincided with shifts in the benthic community as inferred from bird species shifting from primarily bivalve carnivores towards polychaete carnivores (Eriksson, et al. 2010, Van Roomen, et al. 2005) and the drivers of these shifts remain unclear.

Analysis of the stable isotope composition of carbon (δ^13^C) and nitrogen (δ^15^N) in animal tissues are routinely used to identify underlying resource use and trophic relationships within ecosystems (Fry 1988, Minagawa and Wada 1984). Through the application of species specific trophic discrimination factors (TDFs) for carbon and nitrogen (Δ^13^C and Δ^15^N) stepwise isotopic increases that occur between a consumer’s diet and their tissues during metabolism can be accounted for (McCutchan, et al. 2003, Post 2002), allowing for identification of the animal’s trophic level. Accounting for isotopic changes across trophic levels allows for the application of both simple and complex stable isotope mixing models (SIMMs) to be used to resolve the relative use of food resources by consumers within a food web (Stock, et al. 2018, Fernandes, et al. 2014). For the Dutch part of the Wadden Sea, Christianen, et al. (2017) identified that the majority of the biomass is supported by both pelagic phytoplankton and microphytobenthos. Their study used a two-source mixing model with a single isotopic tracer (δ^13^C) and clearly identified benthic productivity from microphytobenthos (MPB) as an important resource supporting the benthic community. However, they may have overestimated the amount of support due to compression of the MPB endmember through the use of an animal proxy instead of directly measured resources (Post 2002).

Thus far, it has not been possible to assess the trophic structure within the Dutch Wadden Sea benthic community due to the large variability in δ^15^N values observed in multiple benthic species (Christianen, et al. 2015). This work found clear trends of increased δ^15^N values for filter feeders (*Mytilus edulis*) in the eastern Dutch Wadden Sea indicating the potential for either an increased δ^15^N baseline value or trophic structure. Potential causes for this increase could be 1) a missing source of N to the ecosystem (e.g. terrestrial input), 2) widespread increases in the trophic complexity resulting in increased δ^15^N values throughout the benthic food web in certain regions, or 3) increased biogeochemical processing of N substrates resulting in increased variability in δ^15^N values due to locally increased denitrification. Terrestrial inputs have been found to be limited within the Dutch Wadden Sea (Christianen, et al. 2017). Intertidal areas within the German Wadden Sea have been identified as having high rates of denitrification associated with tidal pumping and long exposure times within intertidal sediments (Marchant, et al. 2017, Marchant, et al. 2016, Marchant, et al. 2018). Increased porewater processing of nitrogen may coincide with the unique distribution of sandy sediments and pore sizes due to the parallel flow of tidal waters through the Wadden Sea basin. These physical attributes result in reverse grading of sediments with coarser grains towards the basin edges and finer sediment centrally deposited along with longer exposure times in the east than in West (Compton, et al. 2013, Otto, et al. 1990). This combination of unique physical and biogeochemical settings may contribute to increased underlying δ^15^N values but identifying any use of denitrification-affected N porewater resources by MPB and the food web has remained largely intractable solely using traditional bulk isotope techniques.

MPB thrive in unvegetated sandy sediments where they can contribute significantly to the primary production supporting an ecosystem (Miller, et al. 1996) as benthic diatoms fix carbon and take up nitrogen from the overlying water column and porewaters (Cook, et al. 2007, Riekenberg, et al. 2020, Oakes, et al. 2012). Much of the fixed carbon is excreted as extracellular polymeric substances (EPS) which are sticky, sugar rich substrates that help to stabilize sediment and facilitate diatom motility throughout sand (Goto, et al. 1999, Stal 2010), but also serve as a labile carbon source for heterotrophic bacteria (Taylor, et al. 2013). Diatom motility in sandy sediments is an adaptation to the physiochemical variations that occur within intertidal settings. Diel vertical migration of diatoms coincides with tidal cycles and available sunlight to support favorable conditions for photosynthesis at the surface (Barnett, et al. 2020) and to maximize nutrient availability for cell growth and division in the subsurface (Saburova and Polikarpov 2003). Due to the diel vertical migration of several centimeters and a strong coupling between diatoms and heterotrophic bacteria, MPB actively straddle the boundary between water column and sediment porewaters to maximize resource availability.

Food webs are often described as either green or brown depending on whether consumers are supported by herbivory of primary producers or supported by detrital reworking via the micobial loop (Middelburg 2014, Potapov, et al. 2019). MPB supported food webs blur the line between green (MPB_green_) and brown (MPB_brown_) designations as MPB-derived material that can be used by consumers comes in several forms: 1) newly fixed organic matter from water column nutrients (MPB_green_ throughout), 2) MPB-derived organic matter formed from recycled detrital material by heterotrophs via oxic or anoxic pathways in sediment porewaters, or 3) detrital MPB derived material such as reworked EPS that is still labile within the sediment (MPB_brown_ throughout). Microbial subsidy to MPB of both C and N from heterotrophic bacterial reworking of organic matter in porewaters blurs the line between primary production and detrital resource use. Consumers such as deposit feeders have access to MPB biomass (living or dead) or EPS made using a mixture of resources derived directly from the water column or from heterotrophic processing within porewaters (e.g. denitrification affected N pools). This biogeochemical complexity can make identification of carbon and nitrogen food web support from MPB difficult to identify using bulk isotope techniques.

Application of compound specific stable isotope analysis techniques (e.g. analysis of individual amino acid δ^15^N values) could allow for identification of the trophic structure of the benthic community in the Wadden Sea without requiring measurement of all primary producers and all N sources within the ecosystem. This is because amino acid δ^15^N values provide additional information for each sample analyzed in comparison to traditional bulk analysis due to the different fractionations that occur during metabolic reworking of trophic and source amino acids (O’Connell 2017). Trophic amino acids such as glutamic acid (Glu) fractionate considerably between diet and the consumer resulting in a TDF ranging from 3-3.8‰ in marine mammals and sharks (Ruiz-Cooley, et al. 2021, Whiteman, et al. 2018) to as high as 11‰ in teleost fish fed a diet of low protein quality (McMahon, et al. 2015). δ^15^N values for source amino acids such as phenylalanine (Phe) are relatively preserved as they are processed throughout food webs due to a low and consistent TDF of ∼0 to 0.5‰ (McMahon, et al. 2015, Chikaraishi, et al. 2009). Integration of source amino acid values into trophic level estimates removes effects from baseline N variations across an ecosystem using a single chemical analytical method (Xing, et al. 2020, Vokhshoori, et al. 2019) instead of requiring multiple comparisons and adjustments using resource or primary consumer measurements. The combination of two isotopic tracers (δ^13^C and δ^15^N) and amino acid source and trophic δ^15^N values reveals contribution from particulate organic matter as well as two additional sources, MPB_green_ and MPB_brown_ via primary production and use of detrital resources, respectively. This additional information provided by the difference between trophic and source amino acids has the potential to resolve the trophic structure and further clarify MPB resource use supporting the benthic food web in the Wadden Sea.

In this study, we apply natural abundance stable isotope analysis of amino acid nitrogen to resolve the trophic structure for 28 species from within the Dutch part of the Wadden Sea benthic ecosystem which was previously not possible using solely bulk isotope values. Through the application of this amino acid-based method we further confirmed considerable support from MPB-derived carbon relative to phytoplankton, but less than previously postulated. The application of source amino acid δ^15^N values across the ecosystem identified the widespread nature of the driver causing increased variability in δ^15^N across the ecosystem, which we had identified as potentially being either 1) widespread variability in underlying biogeochemical processes (denitrification) or 2) unique sources such as freshwater inflow or groundwater influx coming from terrestrial sources. Finally, application of a three source mixing model revealed considerable detrital support within a subset of the food web that was using the MPB_brown_ resource.

## Methods

### Study site

The Wadden Sea is a large intertidal ecosystem formed behind a chain of 12 sand barrier islands that stretch from the Netherlands to Denmark (Wolff 1983, Christianen, et al. 2017). It is an highly productive estuarine ecosystem formed of sedimentary tidal flats that receive direct and indirect terrestrial input from the Ems, Weser, Elbe, IJssel, Muse, and Rhine rivers (Wolff 1983, Eriksson, et al. 2010). Extensive long-term sampling of the benthic community has occurred in the Dutch Wadden Sea with spatial coverage extending from the Marsdiep to the Ems River mouth at the border of Germany (Fig.1 and Supplementary Fig. 1).

**Figure 1:**
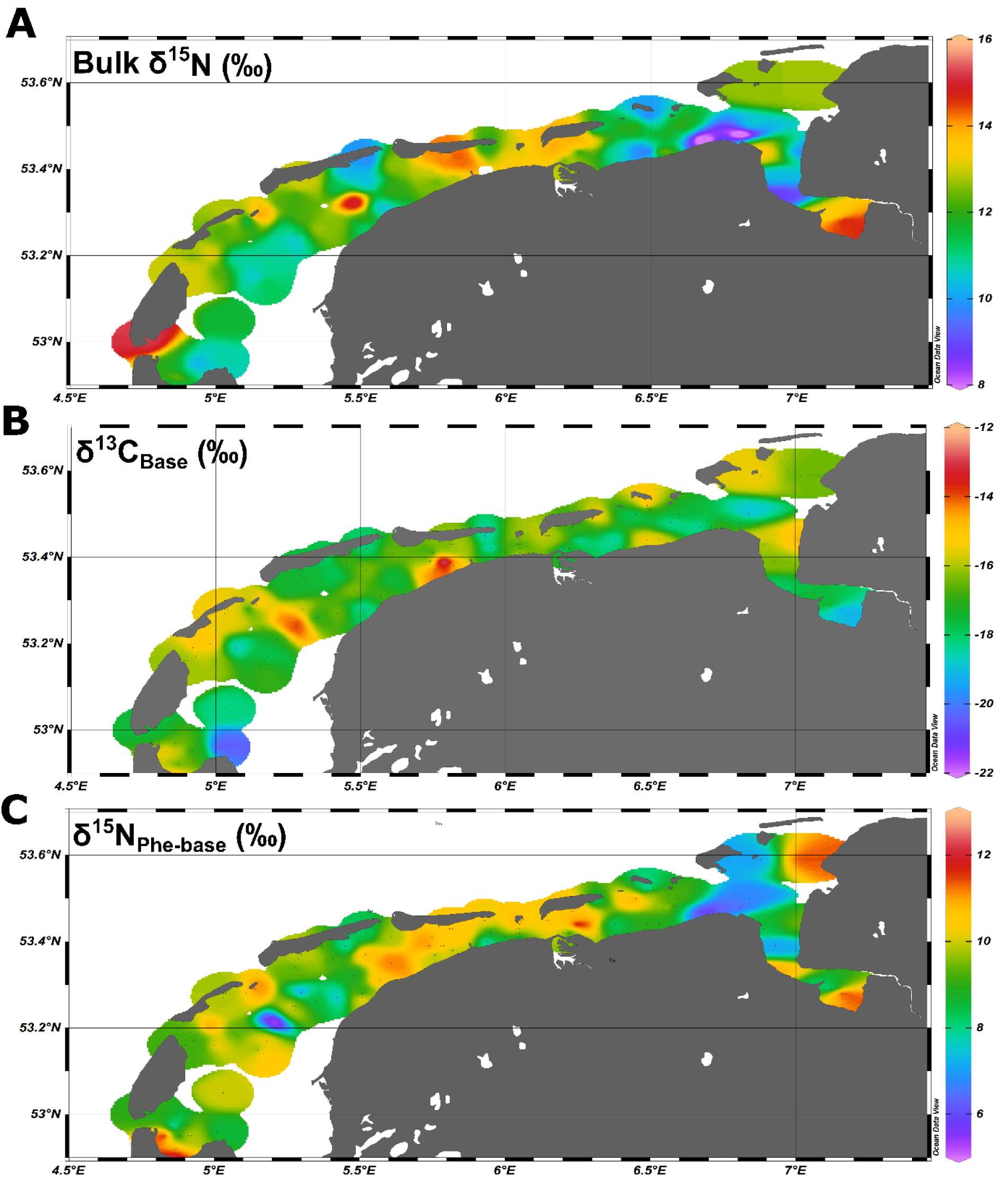
Spatial coverage for A) regionally adjusted bulk δ^15^N values, B) δ^13^C_Base_ values, and C) δ^15^N_Phe-base_ values for the Dutch Wadden Sea.

### Sampling

Sampling for macrozoobenthic species occurred between June and October of 2011-2014 as part of a yearly spatially extensive long-term monitoring campaign within the Dutch Wadden Sea (SIBES, Synoptic Intertidal Benthic Survey). The SIBES program performs core sampling of the intertidal mudflats either by foot or with the use of a rigid hulled inflatable boat, with the aim of comprehensive identification and collection of benthic species in a gridded pattern (500 m separation) throughout the Dutch Wadden Sea with additional random sample points that result in ∼4500 samples per year (Bijleveld, et al. 2012, Compton, et al. 2013). Samples were sieved using 1 mm mesh size and all retained animals were stored frozen after collection. Identification and dissection was done either aboard the ship or in the laboratory. In addition, samples of the filter feeding community (e.g. *Mytilus edulis* and *Balanus crenatus*) were obtained from buoys in the main tidal channels in 2014 to constrain regional N variability throughout the Dutch Wadden Sea. Encrusting communities were scraped from the sides of free-standing anchored buoys and were collected, dissected, and frozen on board (Christianen et al. 2015). Fish were collected from long term monitoring efforts from the NIOZ Fyke. For all sampling campaigns samples of muscle tissue (fish, crustaceans, and bivalves), soft tissue (invertebrates) or whole animals (smaller specimens) were freeze dried, homogenized, and placed in sample vials prior to bulk tissue or amino acid stable isotope analysis. Species were selected for bulk stable isotope analysis based on their contribution to total biomass within the Dutch Wadden Sea ecosystem (Christianen, et al. 2017), with 35 species contributing 99% of the total benthic biomass for the system. From these samples, 28 species were selected for amino acid isotope analysis based on tissue availability (>3 mg), spatial representation across the Dutch Wadden Sea, and adequate replication (>5 individuals) within the data set. Also samples for POM were collected and filtered through combusted GFF filters (Whatman). MPB were filtered onto GFF filters after migration into combusted sand through a 100 μm mesh (Eaton and Moss 1966).

### Isotope analyses

For bulk stable isotope analysis of δ^13^C and δ^15^N values, 0.4 to 2 mg of freeze dried and homogenized animal tissue was loaded into tin or silver capsules depending on whether acidification was required due to the presence of carbonates from shell material. Samples were analyzed with a Flash 2000 elemental analyzer coupled to a Delta V Advantage isotope ratio mass spectrometer (Thermo Scientific, Bremen, Germany). Stable isotope ratios are expressed using the δ notation in units per mil:

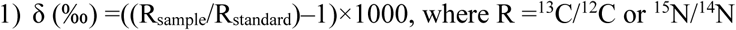

respectively, reported relative to Vienna Pee Dee belemnite and atmospheric N_2._ Laboratory standards of acetanilide, urea, and casein with known δ^13^C and δ^15^N values and known %TOC and %TN values calibrated against NBS-22 and IAEA-N1 were used for calibration with each sample run. Average precision for standards and replicate samples was ±0.1‰ for δ^13^C and 0.2‰ for δ^15^N.

For analysis of δ^15^N values in individual amino acids, 2-5 mg of tissue was hydrolyzed and derivatized to N-pivaloyl/isopropyl (NPiP) derivatives. Analysis occurred using two different gas chromatography combustion isotope ratio mass spectrometers with similar ramp schedules, flow rates and combustion temperatures measured at the NIOZ Royal Netherlands Institute for Sea Research between 2012 and 2019. ∼90% of the samples were analyzed in duplicate with a Trace 1310 gas chromatograph coupled to a Delta V advantage isotope ratio mass spectrometer via an IsoLink II using a modified version of the amino acid preparation and analysis method used by Chikaraishi, et al. (2007) that is described in further detail in Riekenberg, et al. (2020). From this analysis, we report 12 amino acid δ^15^N values including alanine, aspartic acid, Glu, glycine, isoleucine, leucine, lysine, methionine, Phe, serine, threonine, tyrosine, and valine with a precision for samples and standards of <±0.5‰. The remaining 10% of the samples were analyzed in duplicate on an Agilent 6890 gas chromatograph coupled to a Delta V advantage isotope ratio mass spectrometer via a combustion III interface using the method presented in Svensson, et al. (2016) reporting 5 amino acids (alanine, glycine, norleucine, Glu, and Phe) with an average precision for standards and samples of ±1‰. We therefore present only Glu and Phe for the 338 amino acid analyses presented here due to the reduced precision and reduced number of reported amino acids present in 10% of the data set.

### Trophic level calculations

Trophic levels were estimated using single trophic and source amino acids (Glu and Phe, respectively) with the equation presented in Chikaraishi, et al. (2009) modified with an ecosystem specific TDF and β following Bradley, et al. (2015) to account for the range of TDFs present from primary producers to tertiary consumers within the benthic community.

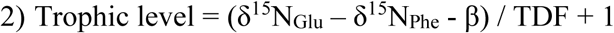

where TDF, the stepwise increase in δ^15^N value between a consumer and their diet is 4.9‰ and β, the difference between Glu and Phe in the ecosystem’s primary producers is 4.3‰. These values are obtained from the slope and intercept of the linear relationship between the difference between δ^15^N values for Glu and Phe and trophic position as determined by stomach content and ecological observation. This relationship was then adjusted to place primary producers at the intercept of 0 using a re-arranged version of equation 1 as further described in (Bradley, et al. 2015):

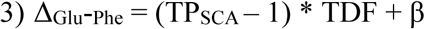

where TP_SCA_ is the trophic level as indicated by stomach content analysis or ecological determination of direct feeding and was determined for each species using FishBase for fish species (Froese and Pauly 2000), literature determinations for invertebrates (Christianen, et al. 2017, Christianen, et al. 2015, Borst, et al. 2018), and interpolation of centroid placement for individual species relative to ecosystem trends (Table 1).

**Table 1:** δ^15^N values for bulk material, glutamic acid, and phenylalanine along with δ^13^C values, trophic position from stomach content and trophic position calculated from amino acids (AAs) for the subset of 25 consumers and 3 macroalgae species examined in this study sampled from the Dutch Wadden Sea benthic food web during SIBES and Waddensleutels campaigns spanning from 2011 to 2014 and in the NIOZ Fyke during routine monitoring.

| Species name | Common name | n | $\delta^{15}\text{N}$ (‰) | SE | $\delta^{13}\text{C}$ (‰) | SE | Trophic Position<br>Stomach Content | Trophic Level From<br>AAs | SE | Glutamic acid<br>$\delta^{15}\text{N}$ (‰) | SE | Phenylalanine<br>$\delta^{15}\text{N}$ (‰) | SE | Glu-Phe<br>$\delta^{15}\text{N}$ (‰) | SE |
| --- | --- | --- | --- | --- | --- | --- | --- | --- | --- | --- | --- | --- | --- | --- | --- |
| Arenicola marina | Lugworm | 14 | 12.8 | 0.4 | -16.0 | 0.5 | 1.9 | 2.1 | 0.1 | 21.0 | 0.7 | 11.2 | 0.5 | 9.8 | 0.4 |
| Balanus crenatus | Acorn barnacle | 13 | 12.5 | 0.2 | -17.4 | 0.4 | 1.9 | 2.4 | 0.2 | 20.7 | 1.0 | 9.3 | 0.4 | 10.9 | 0.8 |
| Carcinus maenas | Green crab | 18 | 14.0 | 0.3 | -15.8 | 0.2 | 3 | 3.2 | 0.1 | 24.2 | 0.7 | 8.9 | 0.3 | 14.5 | 0.6 |
| Cerastoderma edule | Common cockle | 21 | 11.0 | 0.4 | -18.8 | 0.1 | 1.8 | 2.2 | 0.1 | 20.1 | 0.6 | 9.8 | 0.5 | 10.0 | 0.5 |
| Chrysaora hysoscella | Compass jellyfish | 8 | 13.4 | 0.3 | -19.2 | 0.5 | 3.4 | 3.5 | 0.1 | 26.7 | 0.3 | 10.4 | 0.6 | 15.8 | 0.5 |
| Clupea harengus | Herring | 10 | 14.6 | 0.2 | -18.0 | 0.3 | 3.4 | 3.4 | 0.0 | 23.6 | 0.7 | 7.4 | 0.7 | 15.3 | 0.2 |
| Crangon crangon | Shrimp | 17 | 13.6 | 0.3 | -14.9 | 0.3 | 3.2 | 3.5 | 0.1 | 26.2 | 0.4 | 9.7 | 0.4 | 15.7 | 0.5 |
| Magallan gigas | Pacific oyster | 13 | 12.3 | 0.3 | -17.9 | 0.2 | 1.8 | 2.2 | 0.1 | 20.0 | 0.6 | 9.6 | 0.3 | 10.2 | 0.6 |
| Crepidula fornicata | Slipper limpet | 8 | 10.5 | 0.3 | -17.4 | 0.1 | 2 | 1.7 | 0.1 | 17.1 | 0.3 | 9.0 | 0.5 | 7.9 | 0.3 |
| Dicentrarchus labrax | European bass | 10 | 15.9 | 0.5 | -15.9 | 0.5 | 3.5 | 4.0 | 0.2 | 27.8 | 0.6 | 8.3 | 0.3 | 17.9 | 0.7 |
| Ensis leii | Razor clam | 9 | 9.9 | 0.4 | -17.8 | 0.3 | 2 | 1.2 | 0.1 | 14.0 | 0.3 | 8.3 | 0.4 | 5.5 | 0.3 |
| Gobius sp | Goby | 8 | 14.6 | 0.3 | -14.9 | 0.5 | 3.2 | 3.6 | 0.1 | 24.7 | 0.4 | 8.8 | 0.3 | 15.9 | 0.4 |
| Hediste diversicolor | Ragworm | 17 | 12.3 | 0.4 | -16.5 | 0.3 | 2.6 | 2.6 | 0.1 | 20.1 | 0.7 | 7.8 | 0.3 | 11.6 | 0.6 |
| Peringia ulvae | Cone mudsnail | 10 | 9.2 | 0.3 | -15.1 | 0.8 | 2 | 1.4 | 0.1 | 15.3 | 0.5 | 8.2 | 0.7 | 6.7 | 0.5 |
| Lanice conchilega | Sand mason worm | 11 | 11.1 | 0.5 | -18.1 | 0.3 | 1.8 | 2.2 | 0.1 | 19.3 | 0.7 | 9.1 | 0.5 | 9.9 | 0.6 |
| Liocarcinus holsatus | Flying crab | 7 | 13.7 | 0.4 | -18.1 | 0.5 | 3 | 3.3 | 0.1 | 26.5 | 0.4 | 11.7 | 0.6 | 14.8 | 0.3 |
| Littorina littorea | Periwinkle | 17 | 11.5 | 0.3 | -14.5 | 0.4 | 2.3 | 2.3 | 0.1 | 20.7 | 0.4 | 9.8 | 0.2 | 10.6 | 0.4 |
| Macoma balthica | Baltic clam | 20 | 11.9 | 0.3 | -16.2 | 0.2 | 1.8 | 1.7 | 0.1 | 18.5 | 0.6 | 10.1 | 0.4 | 7.8 | 0.5 |
| Mytilus edulis | Blue mussel | 30 | 11.1 | 0.3 | -18.6 | 0.2 | 2 | 1.7 | 0.1 | 17.2 | 0.3 | 8.7 | 0.3 | 8.1 | 0.3 |
| Osmerus eperlanus | European smelt | 8 | 16.4 | 0.4 | -17.2 | 1.0 | 3.5 | 3.8 | 0.1 | 27.0 | 0.5 | 9.4 | 0.4 | 17.1 | 0.3 |
| Platichthys flesus | European flounder | 8 | 15.1 | 0.5 | -14.5 | 0.4 | 3.5 | 3.4 | 0.1 | 25.7 | 0.7 | 9.9 | 0.3 | 15.3 | 0.4 |
| Pleuronectes platessa | European plaice | 10 | 13.4 | 0.3 | -15.6 | 0.3 | 3.2 | 3.2 | 0.1 | 22.7 | 0.5 | 7.4 | 0.5 | 14.2 | 0.4 |
| Rhizostoma pulmo | Barrel jellyfish | 7 | 13.0 | 0.3 | -20.5 | 0.9 | 3 | 2.9 | 0.2 | 23.5 | 0.6 | 10.1 | 0.3 | 13.1 | 0.7 |
| Solea solea | Sole | 11 | 14.8 | 0.2 | -16.8 | 0.3 | 3.2 | 3.0 | 0.1 | 25.5 | 0.6 | 11.4 | 0.5 | 13.6 | 0.6 |
| Zoarces viviparus | Eelpout | 8 | 15.9 | 0.2 | -15.0 | 0.2 | 3.5 | 3.8 | 0.1 | 27.6 | 0.5 | 10.4 | 0.3 | 16.8 | 0.4 |
| Ceramium rubrum | Red horn weed | 8 | 11.6 | 0.3 | -18.4 | 0.9 | 1.5 | 1.7 | 0.1 | 14.5 | 0.7 | 6.6 | 0.8 | 7.7 | 0.3 |
| Fucus vesiculosus | Bladderwrack | 8 | 8.0 | 0.9 | -15.3 | 0.6 | 1 | 0.3 | 0.1 | 12.1 | 0.8 | 10.3 | 0.8 | 1.8 | 0.3 |
| Ulva spp | Sea lettuce | 8 | 11.1 | 0.7 | -13.5 | 0.6 | 1 | 0.8 | 0.1 | 15.7 | 0.6 | 11.5 | 0.4 | 4.1 | 0.4 |

Adjustment to account for the small trophic increase in δ^15^N values of the source AA phenylalanine to establish baseline δ^15^N values for individual species was calculated as:

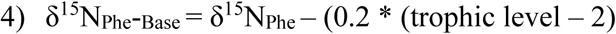

which accounts for the small increase observed in the δ^15^N_Phe_ (‰) value that occurs during metabolism (Chikaraishi, et al. 2009) and trophic level is the trophic level estimate for each individual species. Correction for the trophic increase in bulk δ^13^C values was calculated as:

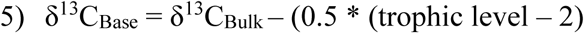

where 0.5 is the increase observed in bulk δ^13^C values between diet and consumer. This value falls between the TDF values observed for bulk δ^13^C for whole tissue from consumers (0.3 ± 0.1‰) and consumer muscle tissues (1.3 ± 0.3‰; (McCutchan, et al. 2003). Due to the wide range of animals included in this analysis ranging from invertebrate primary consumers to teleost tertiary consumers, we have found it appropriate to use a single intermediate TDF value for bulk δ^13^C for all consumers across the trophic structure.

### Data analysis

Statistical analyses were performed in R (v 4.0.3) using RStudio (v 1.3.1056), OriginLab 2020b, or in Ocean Data View (5.6.0, Schlitzer 2022). Mixing models were performed using Food Reconstruction Using Isotope Transferred Signals (FRUITS v 2.1) and the MixSIAR package in R (Stock, et al. 2018). We first used FRUITS to model the species means against measured values for end members (Table 1) for two sources (POM and MPB average) and then three sources (POM, MPB_green_, and MPB_brown_) with SD values set to 0.2‰ for δ^13^C and 0.5‰ for δ^15^N reflecting global measurement errors. Models in FRUITS were run using 50000 updates, a burn-in of 10000 and a minimum uncertainty of 0.001. We then proceeded to modeling individuals within species groupings in MixSIAR using end member values for δ^15^N_Phe_ that allowed for resolution of individuals within the mixing envelope between end members (Table 1b) with minimal exclusion of consumer data points (4%, n=302). Models in MixSIAR were run using the ‘long’ setting, where 3 Markov chain Monte Carlo algorithms with a length of 300,000 run with a burn-in of 200,000. Results are presented from the species averages along with correlation indices comparing the two modeling approaches. Additional details about model fit are provided in the Supplemental Materials.

## Results

Bulk δ^13^C and δ^15^N values and standard errors (SE) for the 28 species included in this study are listed in Table 1 and bulk δ^13^C and δ^15^N values and standard errors (SE) for resources identified as supporting the ecosystem are listed in Table 2a. We confirmed that bulk δ^15^N values were increased for samples taken from the Ems-Dollard region (eastern Dutch Wadden Sea) in comparison to other regions in the Dutch Wadden Sea and have now adjusted for this regionally observed increase in bulk δ^15^N values for filter feeders (−2.3‰; One-way ANOVA, F_2,43_=6.8, *p*=0.003; Supplemental Table 1). Adjustment of δ^15^N values was identified as necessary in both *M. edulis* and *B. crenatus* from samples pooled into regional categories within the Wadden Sea (West <5.5°E, East >5.5°E, and Ems-Dollard associated with the mouth of the river Ems). Despite application of a regional adjustment, ranges for bulk δ^13^C and δ^15^N values remained large for consumers (Fig. 1a and 1b) as well as for particulate organic matter (POM) and MPB sampled from across the Dutch Wadden Sea (POM -15.4 to -21.7‰; 13.7 to 5.7‰ and MPB -10.4‰ to -16.2‰; 13.6 to 6.5‰).

**Table 2:** A) Bulk δ^15^N and δ^13^C values for the end member values supporting the Dutch Wadden Sea benthic food web used for two and three source mixing models on species means. B) Bulk δ^15^N and δ^13^C values for the end member values supporting the Dutch Wadden Sea benthic food web used for two and three source mixing models on individuals within each species.

**A**
| End Members | n | $\delta^{15}\text{N}$ (‰) | SE | $\delta^{13}\text{C}$ (‰) | SE |
| --- | --- | --- | --- | --- | --- |
| Particulate organic matter | 101 | 8.6 | 0.1 | -21.2 | 0.1 |
| Microphybenthos average | 95 | 9.2 | 0.2 | -13.4 | 0.2 |
| Microphybenthos green | 20 | 7.3 | 0.2 | -14.2 | 0.3 |
| Microphybenthos brown | 25 | 11.3 | 0.3 | -13.4 | 0.2 |

**B**
| End Members | $\delta^{15}\text{N}$ (‰) | SE | $\delta^{13}\text{C}$ (‰) | SE |
| --- | --- | --- | --- | --- |
| Particulate organic matter | 8.6 | 0.1 | -21.2 | 0.1 |
| Microphybenthos average | 9.2 | 0.2 | -13.4 | 0.2 |
| Microphybenthos green | 5.0 |  | -14.2 | 0.3 |
| Microphybenthos brown | 13.0 |  | -13.4 | 0.2 |

Increased δ^15^N_Phe_ values were observed for filter feeders sampled from Ems-Dollard in comparison to rest of the Dutch Wadden Sea and values were adjusted in a similar manner as the bulk δ^15^N values (−2.1‰; One-way ANOVA, F_2,43_=5.7, *p*=0.006; Supplemental Table 1). Values for δ^15^N_Phe_ ranged from 6.6‰ observed for *Ciramium rubrum* to 11.7‰ for *Liocarcinus holsatus* and mean values for species correspondingly increased with larger Glu-Phe differences reflecting the minor trophic increases expected for source amino acids (Table 1). δ^15^N_Glu_ values ranged from 12.1‰ for *Fucus vesiculosus* to 27.8‰ for *Dicentrarchus labrax* and were higher than δ^15^N_Phe_ values as expected for the larger fractionation associated with trophic amino acids. δ^15^N values for Glu and Phe were correlated across trophic levels (Fig.2; TL1 R^2^= 0.26 n=24, *p*<0.01; TL 2 R^2^=0.37 n=166, *p*<0.001; TL 3 R^2^=0.39 n=96, *p*<0.002; TL 4 R^2^=0.14 n=34, *p*=0.03), but the relationships were weaker for both TL 1 and TL 4 when compared to TL 2 and TL 3. The relationship between Δ_Glu_-_Phe_ δ^15^N values and TP_SCA_ -1 had a strong correlation (Fig. 3; n=337, R^2^ =0.7, *p*<0.001) with a slope of 4.9±0.3‰ and an intercept of 4.3±0.2‰ that reflect TDF and β, respectively, for the Dutch Wadden Sea benthic food web. These values for TDF and β are different than the canonical values of 7.6‰ and 3.4‰ found by (Chikaraishi, et al. 2009) and resulted in a slope of 0.99 for TL versus TP_SCA_ while using the canonical TDF and β resulted in a slope of 0.64 (both R^2^=0.72).

**Figure 2:**
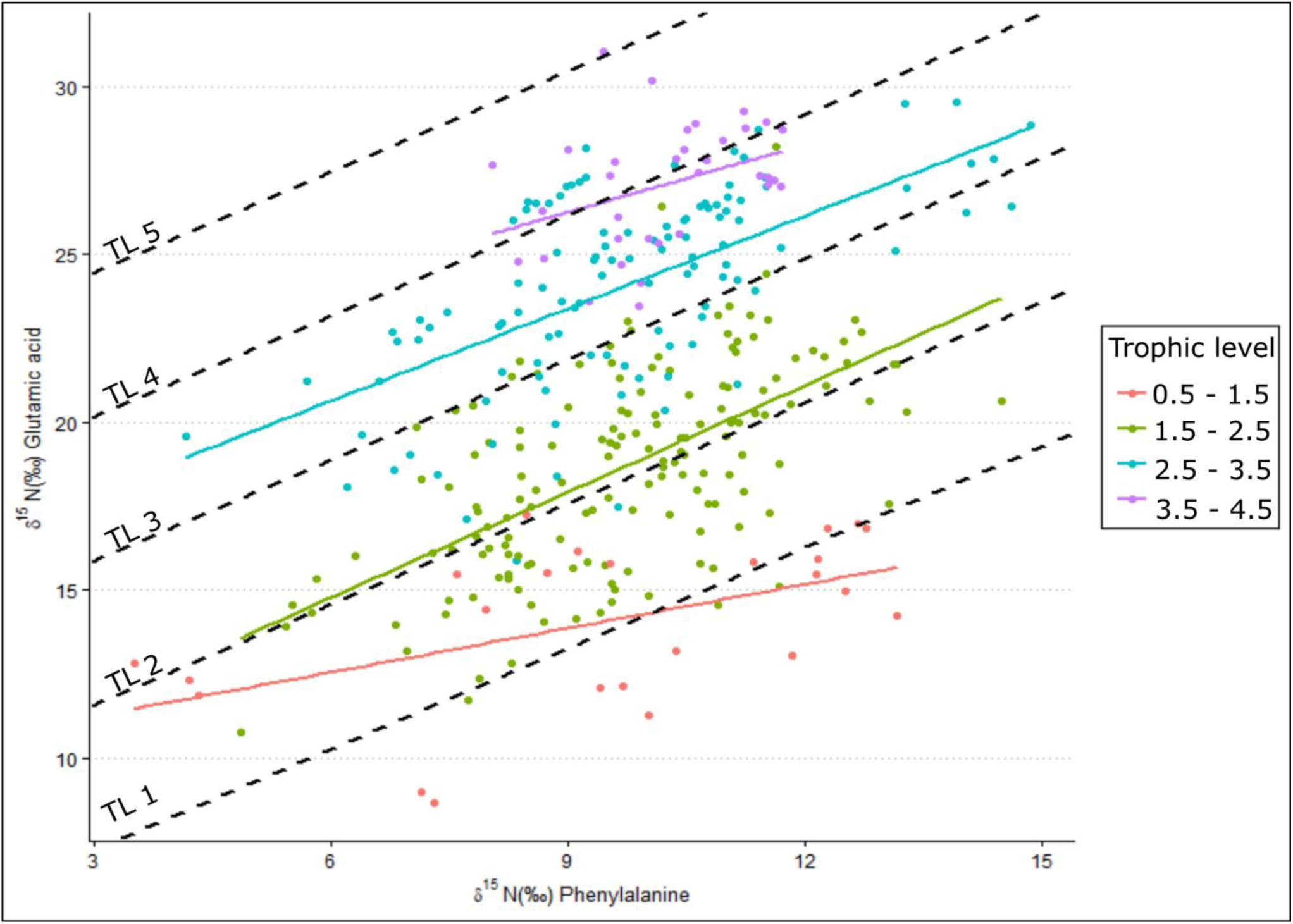
δ^15^N values for the amino acids glutamic acid and phenylalanine measured for species across different trophic levels. Dashed lines are expected linear relationships for trophic levels 1 to 5 using a trophic discrimination factor of 4.9‰ and a β of 4.3‰, while colored linear regressions are trophic level relationships by grouping trophic levels across species (n=24-166).

**Figure 3:**
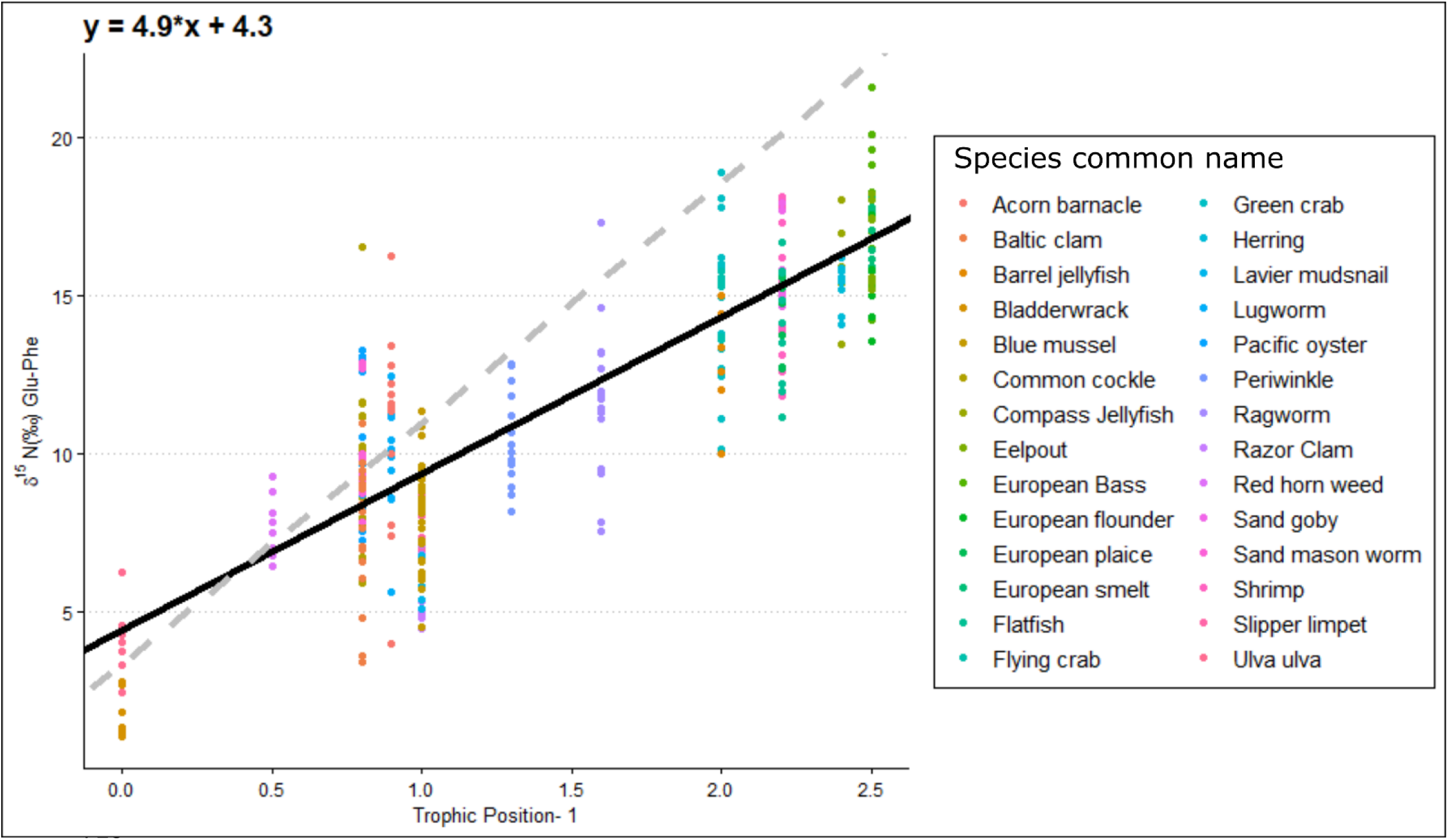
The per mil difference in δ^15^N value between glutamic acid and phenylalanine in relation to trophic levels for individual species minus one level to account for primary producers. The black line is the fitted linear regression across all species (r^2^=0.7, n= 337) with the equation presented at the top left of figure. The dashed grey line is the linear trophic level relationship using a slope of 7.6‰ and a β of 3.4‰ as presented in Chikaraishi et al. (2009).

Calculated trophic levels for primary producers ranged from 0.3 for *Fucus vesiculosus* to 1.7 for *C. rubrum* (Fig. 4). Primary consumers ranged from 1.2 for *Ensis leii* to 2.3 for *Littorina littorea*, secondary consumers from 2.6 for *Hediste diversicolor* to 3.5 for *Crangon crangon*, and tertiary consumers from 3.6 for *Gobius sp*. to 4 for *D. labrax* (Fig. 3). No amino acid measurements were possible for POM or MPB as part of this study as the sample weights remaining after the initial bulk analysis between 2012 and 2015 were below detection limits for this analysis. Consumer δ^13^C values fell between the measured values for POM and MPB (−21.2‰ and -13.4‰). We therefore used the trophic level estimates calculated using system-specific TDF and β to account for fractionation while estimating values for δ^13^C_Base_ and δ^15^N_Phe-base_.

**Figure 4:**
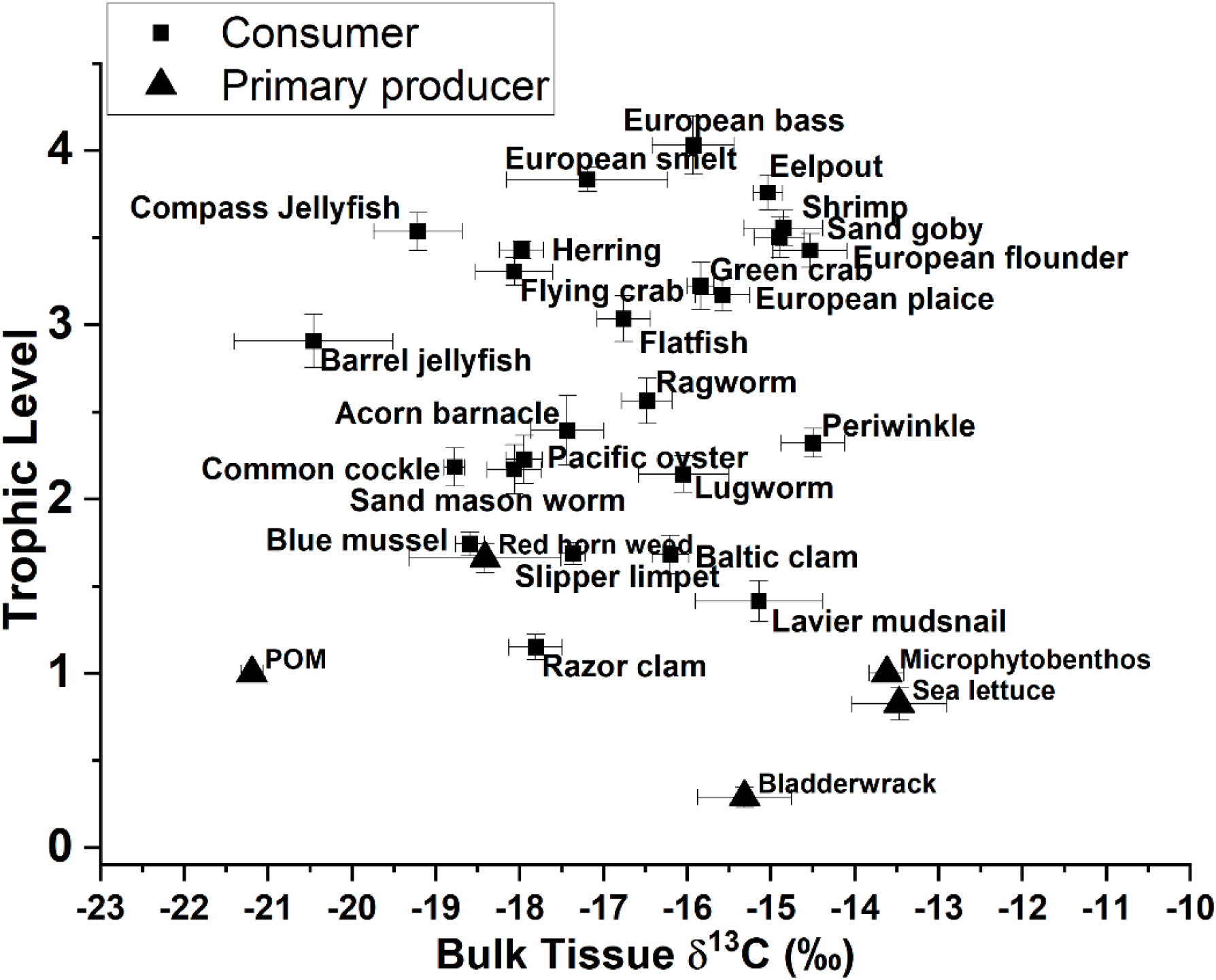
Trophic level estimates from amino acid δ^15^N values for glutamic acid and phenylalanine versus bulk δ^13^C values for both consumers and primary producers.

We then used δ^13^C_Base_ values from POM and MPB (Table 2) and each consumer to calculate dietary contributions for species averages (Fig. 5a) and for all individuals within each species. Dietary contributions from these 2 source mixing models are presented in Supplementary Tables 2a and indicated use of POM (>0.5) for 11 of the 25 species examined with the highest contribution being observed for *Rhizostoma pulmo* (0.86±0.09 and 0.76±0.04). To further investigate the use of MPB within the ecosystem, we used δ^13^C_Base_ and δ^15^N_Phe-Base_ values from POM, MPB_green_, a proxy of freshly fixed MPB-derived material, and MPB_brown_, a proxy of MPB-derived material using reworked N and OM associated with heterotrophic reworking of detrital OM and denitrification occurring across the tidal cycle (Table 2b), to calculate dietary contributions from species average values (Fig. 5b). Both MPB_green_ and MPB_brown_ end members are from minimum and maximum bulk δ^15^N values for MPB measured during the sampling campaigns. Similar to the 2 source models, dietary contribution from POM was considerable (>0.5) for 11 of the 25 species with *R. pulmo* having the highest contribution (0.78±0.1). For the two MPB end members, dietary contribution of MPB_green_ was considerable for 6 species with *Pleuronectes platessa* having the highest contribution (0.75±0.1) and dietary contribution from MPB_brown_ was considerable for 5 species with *Zoarces viviparus* having the highest contribution (0.57±0.13). To calculate dietary contributions from individuals, end member values for N were expanded for both MPB-derived end members, with δ^15^N values of 5.0‰ for MPB_green_ and 13.0‰ for MPB_brown_ taken from values observed for δ^15^N_Phe-Base_ for individuals from *Hydrobia ulvae* and *Littorina littorea*. Dietary contributions from POM were >0.5 for 10 species with *Chrysaora hysoscella* having the highest contribution (0.78±0.04), while contributions for MPB_green_ were lower with no species >0.5, but with 4 species having a contribution of >0.4 with *Hediste diversicolor* having the highest contribution (0.47±0.05), and contributions for MPB_brown_ being >0.5 for 6 species with *Arenicola marina* having the highest contribution (0.67±0.05). Both two-source and three-source mixing models indicated good agreement between the use of POM with a correlation between the models using species averages and individual values of 0.88 and 0.89 respectively (R^2^, Fig. 5c).

**Figure 5:**
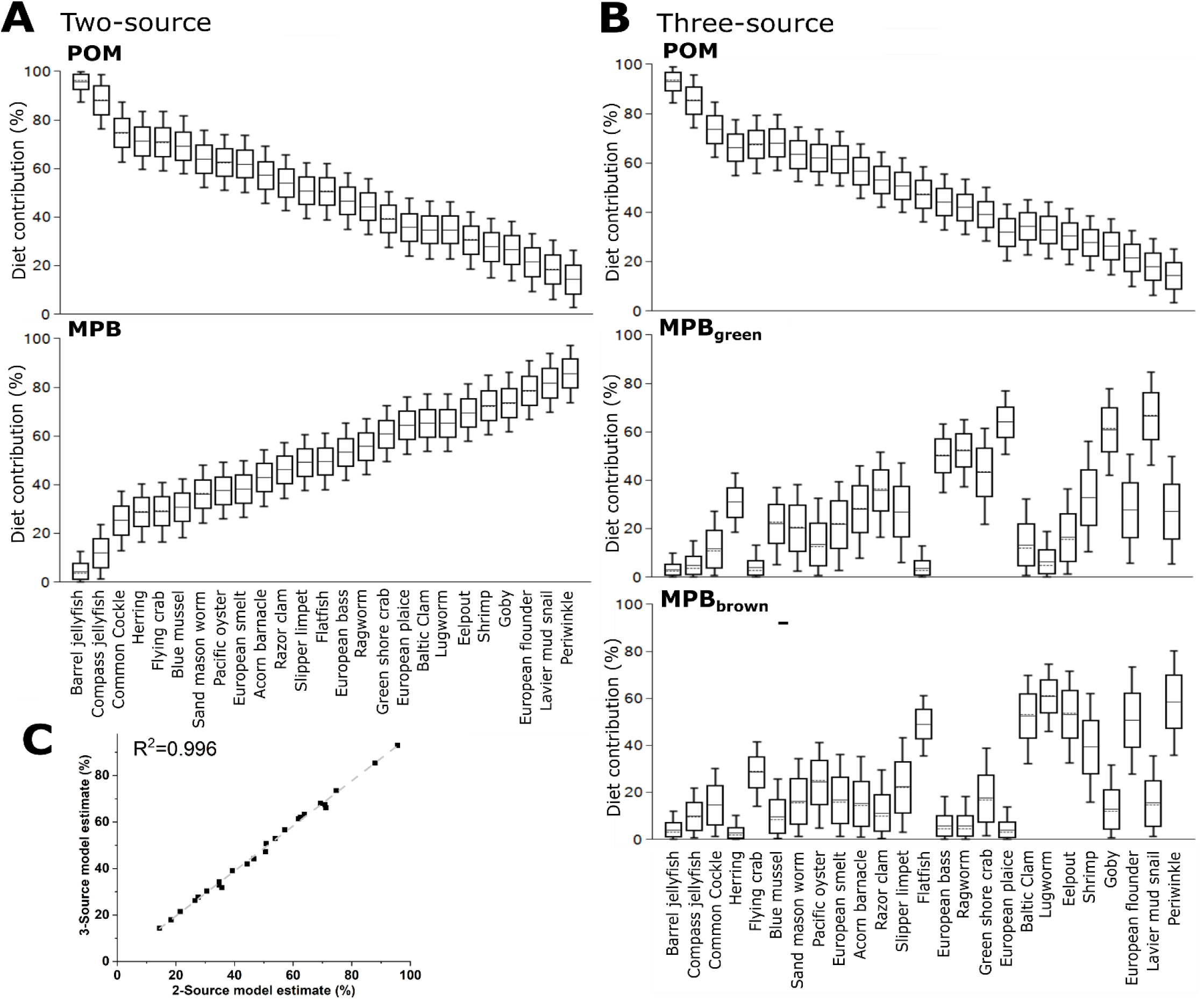
Estimated dietary contributions from primary producers using A) a two source mixing model using only δ^13^C and mean values for each species and B) a three source mixing model using both δ^13^C and δ^15^N_Phe-Base_ and mean values for each species and C) the correlations between two and 3 source mixing models when applied to means values and individuals for each species. Trophic structure has been accounted for in both models using trophic level estimates from amino acid analysis of δ^15^N.

## Discussion

In the Dutch Wadden Sea, we found that using amino acid δ^15^N values allowed for more detailed and reliable identification of the trophic structure of the benthic foodweb. This improvement allowed for the identification of a subset of species that rely on microbially reworked materials supported through active tidal pumping. Across all four simple and complex stable isotope mixing models (SIMMs) mean consumer reliance on MPB was 0.55 to 0.6, but fell short of the 0.7 previously observed (Christianen, et al. 2017). This smaller contribution from MPB is likely due to the combined use of δ^13^C and δ^15^N values to assess resource contributions instead of using consumer integrators as a proxy for MPB resource values (Post 2002). Additionally, analysis of individual amino acid δ^15^N values allowed for trophic adjustment of δ^13^C and δ^15^N values to account for trophic discrimination for the 25 benthic species prior to analysis with SIMMs. Through the application of source amino acids, we were able to construct a trophic structure within the Dutch Wadden Sea despite the considerable underlying variability present in bulk δ^15^N values across the area (Fig. 1a). This variability was likely caused by widespread biogeochemical processing unique to regions of intense tidal flushing within intertidal sands (e.g. denitrification). Analysis of trophic and source amino acid δ^15^N values allowed for the development and application of a system specific TDF and β to determine the trophic structure of the Wadden Sea benthic food web that was previously impossible using solely bulk isotopic techniques.

We confirm that MPB is a major source of productivity that widely supports the food web in the Dutch Wadden Sea (Christianen et al. 2017). The majority of consumers examined here (21 out of 25 species) have δ^13^C values higher than -18‰, indicating reliance on a resource with a higher value than POM (−21.2±0.2‰; n=101). MPB (−13.4±0.2‰; n=95) is a widely available resource across the Dutch Wadden Sea (Fig.1b) with variable δ^13^C values indicating use of both dissolved inorganic carbon (DIC) from the overlying water column and remineralized organic matter from porewater DIC. Due to the sandy nature of the sediment and extensive tidal pumping that occurs in the basin, benthic associated MPB, primarily pennate diatoms (Scholz and Liebezeit 2012) are widely available to consumers during diel inundation and exposure. Other potential candidate resources available to consumers include sediment organic matter, terrestrial input, macroalgae or seagrasses, but are unlikely to play a significant role in supporting production. Sediment organic matter (−21.8±0.3‰; n=117) and terrestrial inputs (−27‰, Middelburg and Herman 2007) and -23‰ (Jung, et al. 2019) have lower δ^13^C values than POM and therefore cannot contribute to explain the higher values widely observed across consumers in the Dutch Wadden Sea. Green macroalgae such as *Ulva* (−13.5±0.6‰, n=8) have a higher value that could meaningfully contribute but are spatially limited due to their need for hard anchoring points such as exposed rocks within the larger landscape of mud and sand. This limited distribution constrains the amount of contribution from this resource as blooms of *Ulva* or so-called green tides (Charlier, et al. 2008) did not occur in the period between 2011 and 2014 in the Dutch Wadden Sea. This makes *Ulva* an unlikely resource to support more than half of the food web. Similarly, seagrasses have previously occurred extensively in the Wadden Sea, but due to their current extremely limited spatial ranges in the German and Danish Wadden Sea (Folmer, et al. 2016) are unlikely to be a large source of productivity supporting consumers in the Dutch Wadden Sea.

Further examination of MPB as a resource reveals substantial variability in δ^15^N values ranging from 7.3‰ to 11.3‰ when grouping the minimum and maximum measurements taken from across the study site (Table 2a; MPB_green_ and MPB_brown_). This variability could potentially reflect input from an unaccounted resource or a biogeochemical process that is occurring across the basin. Although freshwater input from terrestrial sources has a δ^15^N of 10-14‰ (Jung, et al. 2019), the limited inputs to the Wadden Sea and δ^13^C value of -23‰ indicate at best a regionally confined contribution to the ecosystem. Another possible unaccounted source includes submarine groundwater discharge, which due to the relatively porous nature of the sandy sediment in the Wadden Sea basin could contribute reworked N with higher δ^15^N values to the benthos. Previous work has identified freshwater groundwater discharge as a minor input (Santos, et al. 2015) that is unlikely to have a basin wide impact. However, this work did confirm considerable porewater exchange resulting in efflux of waters with higher total dissolved nitrogen and lower dissolved oxygen saturation due to tidal pumping. Porewater exchange across the sediment surface resulted in porewater total dissolved nitrogen that decreased with depth that is expected with efflux into the overlying water column. Tidal pumping through permeable sands (Marchant, et al. 2018) is a potentially region-wide process providing nutrients that have undergone considerable denitrification that may provide substrate pools in porewaters with increased δ^15^N values that then support MPB production within the Wadden Sea. Increased δ^15^N values are most apparent in the Ems-Dollard region and required adjustment for both bulk δ^15^N and δ^15^N_Phe-Base_ values (Supplemental Table 1). Despite this adjustment within the wider data set, there remains considerable variability in δ^15^N values (Fig. 1a) across the basin that has made identifying a trophic structure using bulk δ^15^N values alone difficult without a method to integrate underlying shifts in baseline δ^15^N values across the Dutch Wadden Sea.

The TDF value from analysis of the difference between Glu and Phe throughout the food web is lower in this study than the most often applied canonical value (4.9‰ compared to 7.6‰; Chikaraishi, et al. 2009) as has been previously observed in multiple studies (McMahon, et al. 2015, Lemons, et al. 2020, Hebert, et al. 2016, Nuche-Pascual, et al. 2021). The lower TDF value reflects an ecosystem wide application to species ranging from macroalgae to teleosts and represents a compromise between application of multiple species-specific TDFs as determined by feeding studies (28×, 1 per species) or feeding group dependent strategies based on the relative diet vs tissue quality comparisons (McMahon, et al. 2015). By using wild caught animal across an entire ecosystem to derive a system wide TDF, we relied on measured variations in the difference between Glu and Phe between individuals in each species sampling to estimate the TDF value that was then applied to estimate the trophic structure within the ecosystem. To this end, we combined trophic position estimates of teleosts from FishBase (Froese and Pauly 2000), literature values for trophic position of invertebrate species (Christianen, et al. 2015) and measured δ^15^N_Glu-Phe_ values for primary producers to ensure representation across the food web and found a TDF that is broadly comparable to one developed using solely marine teleosts (5.7‰ Bradley, et al. 2015). The resulting system wide TDF represents a compromise that falls between two most often applied approaches for assessing trophic levels 1) using a universally applied canonical TDF (Chikaraishi, et al. 2009, Kato, et al. 2021, Vokhshoori, et al. 2021) and 2) using individual TDFs from either each species or feeding groups from values obtained from controlled feeding studies of similar species (McMahon and McCarthy 2016, Bode, et al. 2021, Le-Alvarado, et al. 2021). Application of a system-specific TDF was possible here due to the relatively large number of replicates for each species examined and having a large number of species for which no controlled feeding studies have been performed using amino acids (e.g. a wide range of benthic invertebrates), especially considering the previously identified specialization on MPB throughout this intertidal food web (Christianen et al 2017).

Application of δ^15^N values from amino acids allows for the integration of underlying variability in N sources when characterizing the trophic relationships within the benthic community. The large range observed for δ^15^N_Phe_-_Base_ (Fig. 1c) in consumers further confirms that use of N with a high δ^15^N value is not regionally confined and occurs across the basin. In contrast, variability in the δ^13^C_Base_ values is more regionally confined, likely indicating the widespread use of MPB-derived carbon that is more consistently available across the basin and therefore less variable. These contrasting patterns and wide range of δ^15^N values observed across the MPB sampling support splitting of the MPB end member into two distinct resources: 1) MPB_green_ which reflects newly fixed organic matter from MPB supported by dissolved inorganic C (DIC) and N substrates from the overlying water column and 2) MPB_brown_ which reflects newly fixed organic matter from MPB supported from bacterially reworked C and N substrates provided from porewater associated materials (Table 2a). Furthermore, δ^15^N_Phe-Base_ values for a subset of the measured individuals of *Peringia ulvae* (5.6±0.2‰; n=3 vs 8.2±0.7‰; n=10) indicated individual specialization on a specific resource in a species previously used as an end member proxy for MPB (Christianen, et al. 2017) while *Littorina littorea* (9.8±0.2‰; n=17; table 1) had a substantially higher δ^15^N_Phe-Base_ value despite similar dietary contribution for MPB between these two species indicated by the two source mixing model (Fig. 5A; 71% and 82%, respectively). A 5‰ difference between source amino acid δ^15^N values between two primary consumers using similar amounts of MPB derived carbon further confirms the substantial variation in MPB-δ^15^N values due to availability of porewaters that have been impacted by denitrification. This difference warranted further examination using three-source mixing models with two distinct MPB sources after adjustment for trophic position indicated using the ecosystem-dependent TDF from amino acid δ^15^N values.

Application of the three source mixing models (Fig. 5b) reveal separation between species groups that use the MPB_green_ and the MPB_brown_ resources. Species using MPB_green_ include *Carcinus maenas, Gobius spp*., *Pleuronectes platessa*, and *P. ulvae*, indicating reliance on newly fixed MPB-derived organic matter supported from the overlying water column (DIC and available N). Species using MPB_brown_ include *M. baltica, A. marina, L. littorea, Crangon crangon, Z. vivparus, Platicthys flesus* and *Solea solea* indicating reliance on MPB-derived organic matter supported off of microbially reworked N substrates that are readily available due to tidal pumping (Fig. 6). Food webs are often split into herbivory (green) and detrital (brown) pathways depending on whether newly fixed productivity or detrital reworking are the basal resource supporting them (Middelburg 2014, Odum 1969). The separation observed between these two groups of species appears to be along the boundaries of green versus brown food webs (Cordone, et al. 2020, Evans-White and Halvorson 2017), where green food webs are directly supported by newly fixed OM from primary producers and brown food webs have detrital and microbial support through the breakdown, processing, and reuse of organic matter through an efficient microbial loop (Azam, et al. 1983, Fenchel 2008). The re-entry and reuse of detrital products (MPB_brown_) in this system appears to be mediated by MPB and may reflect either 1) support of live MPB using reworked nutrients and porewater DIC or 2) the direct use of reworked dead MPB-derived organic matter by deposit feeders. MPB samples in this study required migration across a permeable filter pressed against the sand so both MPB_green_ and MPB_brown_ end members reflect live sampled MPB. The large variability for both bulk δ^13^C and δ^15^N values found in the MPB end members points to the need for further investigation into how MPB are supported across the tidal cycle as well as better characterization for all primary producers using compound specific techniques (PLFAs, AA-C, and AA-N) within the Wadden Sea. Partitioning support from MPB throughout the food web into green and brown pathways highlights that each individual resource in this shallow system (POM, MPB) is likely have multiple forms due to the shallow nature of the basin as green and brown food webs interact and exchange materials (Krumins, et al. 2013). The various forms of organic matter reflect the stages of processing that have occurred: 1). Direct initial use of phytoplankton (green POM), 2). deposition and partial reworking of phytoplankton that are resuspended and used (brown POM/SOM), 3). direct uptake of water column N and DIC (MPB_green),_ 4). use of reworked nutrients and porewater DIC (MPB_brown_), or 5). direct use of detrital MPB biomass (MPB_brown_).

**Figure 6:**
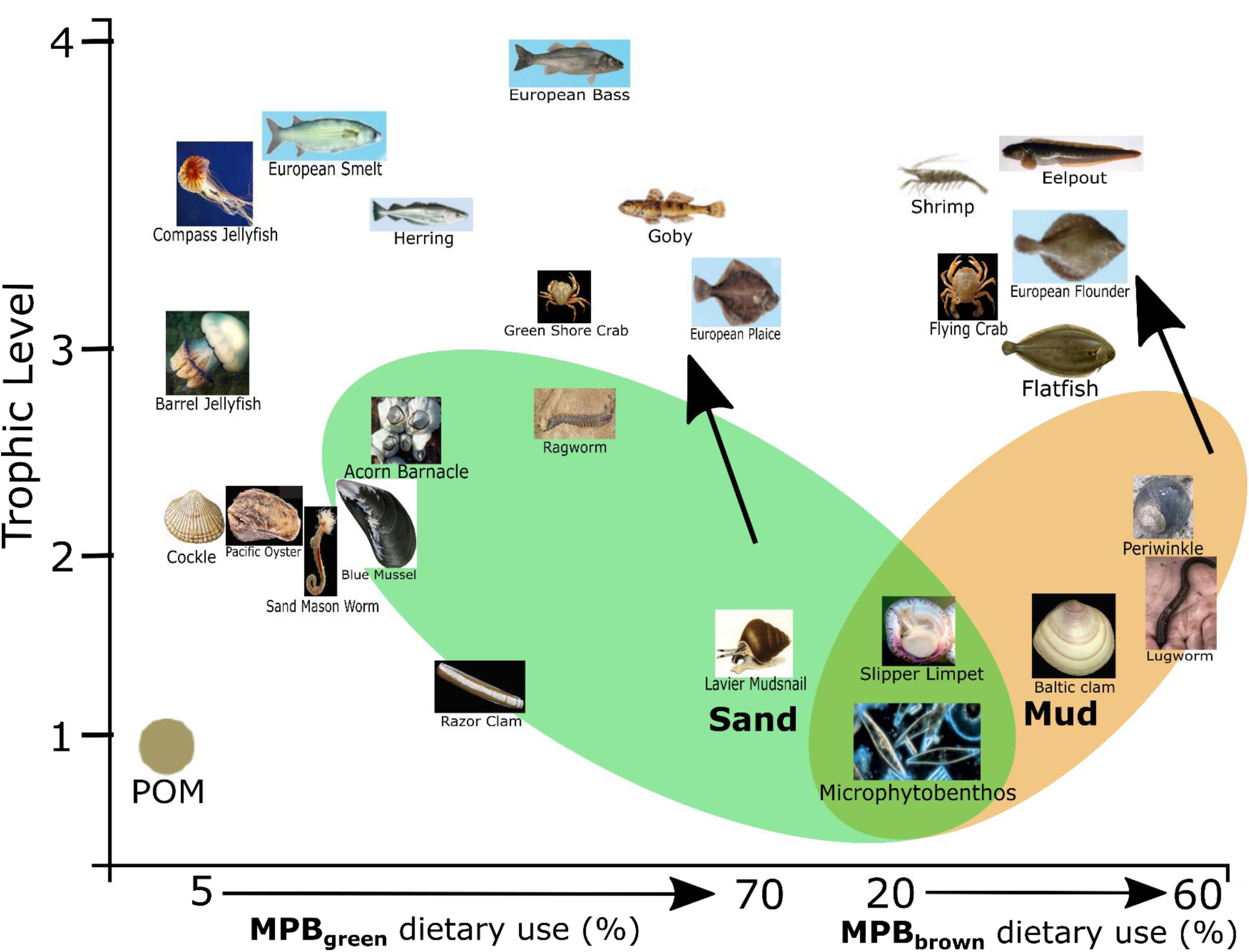
Conceptual diagram indicating increased use of microphytobenthos derived matter from either reworked material (MPB_Brown_) or newly produced material (MPB_green_) as use of particulate organic matter (POM) decreases. Photo credits: Henk Heesen, Ales Kladnik, Francesca Crippa, Hans Hillewaert, Matthisas Buschmann, Sytske Dijksen, and Gordon Taylor.

Due to the diversification of MPB resources, there is support of two distinct food web pathways in the Dutch Wadden Sea for MPB-derived resources as indicated by the three-source mixing models (Figs. 5b and 6): newly fixed production from the overlying water column (MPB_green_) and use of microbially reworked substrates from porewaters (MPB_brown_). This resource diversification likely contributes to the outstanding productivity that has been observed within the Wadden Sea. Specialization on the two different MPB types allows for further feeding niche differentiation beyond the classic model of deposit and surface/particulate feeders through identification of MPB specialist consumers that predominantly rely on either newly fixed or reworked MPB derived organic matter. This specialization likely reduces competition between feeding types in a highly productive ecosystem and reflects efficient use of not only newly fixed organic matter, but also efficient use of reworked OM as either direct uptake or through nutrient and DIC support of MPB through tidal pumping.

## Conclusions

Compound specific analysis of amino acid nitrogen allowed for further identification and adjustment for underlying baseline shifts in δ^15^N values within the Dutch Wadden Sea. The resulting regional adjustments allowed for the development and application of a system-specific TDF that could then successfully be applied to estimate the trophic structure of 28 species within the food web. Further application of stable isotope mixing models confirmed the dominant support of the food web by MPB (2-source models) and identified biogeochemical pathways supporting MPB along separate green and brown pathways (3-source models). Separate support of MPB between newly fixed and reworked organic matter showcases that diatom connectivity with underlying porewaters needs to be further considered in food web studies examining the considerable productivity that occurs within intertidal ecosystems.

## Supporting information

Riekeneberg et al 2022_Supplemental

## Acknowledgements

For sampling the food web, we thank the crew of RV Navicula and the volunteers, staff, and students in the field. We thank Kevin Donkers, Ronald van Bommel, Jort Ossebaar, Monique Verweij, Anchelique Mets, Elisabeth Svensson, and Thomas Leerink (NIOZ) for technical assistance in both bulk and compound specific stable isotope analyses. This study was carried out as part of the project “Waddensleutels” funded by “Waddenfonds” (WF203930). SIBES-monitoring was financially supported by NAM, NWO-ALW (ZKO program) and Royal NIOZ. This study also had support from the UU-NIOZ project “Using Isotopic Indicators” (NZ4543.13).

