## Supplementary material for "Compound specific stable isotope analysis of amino acid nitrogen reveals detrital support of microphytobenthos in the Dutch Wadden Sea benthic food web": Riekeneberg et al 2022_Supplemental

**Supplemental Materials**

**Supplemental Table 1:** Values for filter feeders sampled during the buoy sampling in 2012 across the regions within the Wadden Sea. The regions West and East were split using the dividing line of 5.5° E and Ems-Dollard is the mouth of the river Ems. One-way ANOVAs between the regions indicate a statistical difference for the $\delta^{15}N$ values for bulk material and phenylalanine. The resulting adjustment for the Ems-Dollard region are indicated at the bottom of the top table.

| | | **Bulk $\delta^{15}N$** | | **$\delta^{15}N$ Phenylalanine** | |
|---|---|---|---|---|---|
| Wadden Sea Region | n | Average (‰) | SE | Average (‰) | SE |
| West | 18 | 11.3 | 0.3 | 8.8 | 0.2 |
| East | 19 | 11.8 | 0.2 | 9.5 | 0.4 |
| Eems-Dollard | 7 | 13.8 | 1.2 | 11.2 | 0.8 |
| Adjustment for Eems-Dollard region | | **2.3** | | **2.1** | |

| **One Way ANOVA** | | **One Way ANOVA** | |
|---|---|---|---|
| $F_{2,43}$ | 6.8 | $F_{2,43}$ | 5.7 |
| P | 0.003 | P | 0.006 |
| Tukey's HSD | AAB | Tukey's HSD | AAB |

Supplementary Table 2: Dietary contribution for A) two and B) three source mixing models using average values for species and individual values for species respectively.

**A)**

| | Average values | | | | Individual values | | | | |
|---|---|---|---|---|---|---|---|---|---|
| **Species** | **MPB** | **SD** | **POM** | **SD** | **n** | **MPB** | **SD** | **POM** | **SD** |
| Barrel jellyfish | 0.14 | 0.09 | 0.86 | 0.09 | 23 | 0.24 | 0.04 | 0.76 | 0.04 |
| Common cockle | 0.26 | 0.02 | 0.74 | 0.02 | 19 | 0.27 | 0.02 | 0.73 | 0.02 |
| Blue mussel | 0.27 | 0.03 | 0.73 | 0.03 | 4 | 0.36 | 0.02 | 0.64 | 0.02 |
| Compass jellyfish | 0.29 | 0.07 | 0.71 | 0.07 | 5 | 0.20 | 0.04 | 0.80 | 0.04 |
| Sand mason worm | 0.35 | 0.05 | 0.65 | 0.05 | 15 | 0.46 | 0.04 | 0.54 | 0.04 |
| Pacific oyster | 0.37 | 0.03 | 0.63 | 0.03 | 8 | 0.47 | 0.03 | 0.53 | 0.03 |
| Razor clam | 0.37 | 0.05 | 0.63 | 0.05 | 11 | 0.38 | 0.04 | 0.62 | 0.04 |
| Flying crab | 0.42 | 0.06 | 0.58 | 0.06 | 16 | 0.39 | 0.05 | 0.61 | 0.05 |
| Slipper limpet | 0.43 | 0.03 | 0.57 | 0.03 | 9 | 0.43 | 0.04 | 0.57 | 0.04 |
| Herring | 0.44 | 0.04 | 0.56 | 0.04 | 6 | 0.40 | 0.04 | 0.61 | 0.04 |
| Acorn barnacle | 0.44 | 0.06 | 0.56 | 0.06 | 2 | 0.55 | 0.03 | 0.46 | 0.03 |
| European smelt | 0.57 | 0.13 | 0.43 | 0.13 | 20 | 0.58 | 0.05 | 0.42 | 0.05 |
| Sole | 0.57 | 0.05 | 0.43 | 0.05 | 24 | 0.57 | 0.04 | 0.43 | 0.04 |
| Baltic clam | 0.58 | 0.03 | 0.42 | 0.03 | 18 | 0.58 | 0.03 | 0.42 | 0.03 |
| Ragworm | 0.58 | 0.04 | 0.42 | 0.04 | 13 | 0.64 | 0.03 | 0.36 | 0.03 |
| Lugworm | 0.61 | 0.07 | 0.40 | 0.07 | 1 | 0.73 | 0.04 | 0.27 | 0.04 |
| Green crab | 0.70 | 0.03 | 0.30 | 0.03 | 3 | 0.67 | 0.03 | 0.34 | 0.03 |
| Cone mud snail | 0.71 | 0.10 | 0.29 | 0.10 | 14 | 0.61 | 0.05 | 0.39 | 0.05 |
| European plaice | 0.73 | 0.05 | 0.27 | 0.05 | 22 | 0.67 | 0.04 | 0.34 | 0.04 |
| European bass | 0.74 | 0.07 | 0.26 | 0.07 | 10 | 0.63 | 0.04 | 0.37 | 0.04 |
| Periwinkle | 0.82 | 0.05 | 0.18 | 0.05 | 17 | 0.83 | 0.04 | 0.18 | 0.04 |
| Sand goby | 0.85 | 0.06 | 0.15 | 0.06 | 12 | 0.77 | 0.05 | 0.23 | 0.05 |
| Shrimp | 0.84 | 0.04 | 0.16 | 0.04 | 7 | 0.78 | 0.04 | 0.23 | 0.04 |
| Eelpout | 0.84 | 0.03 | 0.16 | 0.03 | 25 | 0.72 | 0.05 | 0.28 | 0.05 |
| European flounder | 0.88 | 0.06 | 0.12 | 0.06 | 21 | 0.81 | 0.05 | 0.19 | 0.05 |

**B)**

| | Average values | | | | | | Individual values | | | | | |
|---|---|---|---|---|---|---|---|---|---|---|---|---|
| **Species** | **POM** | **SD** | **MPB-sand** | **SD** | **MPB-mud** | **SD** | **n** | **POM** | **SD** | **MPB-sand** | **SD** | **MPB-mud** | **SD** |
| Barrel jellyfish | 0.78 | 0.10 | 0.08 | 0.07 | 0.14 | 0.09 | 23 | 0.75 | 0.04 | 0.07 | 0.04 | 0.18 | 0.05 |
| Common cockle | 0.71 | 0.03 | 0.17 | 0.08 | 0.12 | 0.07 | 19 | 0.71 | 0.02 | 0.16 | 0.04 | 0.13 | 0.04 |
| Blue mussel | 0.73 | 0.03 | 0.10 | 0.07 | 0.17 | 0.07 | 4 | 0.63 | 0.02 | 0.10 | 0.04 | 0.27 | 0.04 |
| Compass jellyfish | 0.67 | 0.07 | 0.09 | 0.08 | 0.23 | 0.09 | 5 | 0.78 | 0.04 | 0.06 | 0.04 | 0.16 | 0.05 |
| Sand mason worm | 0.63 | 0.05 | 0.19 | 0.11 | 0.18 | 0.09 | 15 | 0.52 | 0.04 | 0.20 | 0.07 | 0.28 | 0.06 |
| Pacific oyster | 0.61 | 0.04 | 0.15 | 0.10 | 0.24 | 0.09 | 8 | 0.51 | 0.03 | 0.15 | 0.05 | 0.34 | 0.05 |
| Razor clam | 0.59 | 0.05 | 0.29 | 0.10 | 0.12 | 0.08 | 11 | 0.59 | 0.04 | 0.25 | 0.07 | 0.16 | 0.05 |
| Flying crab | 0.53 | 0.06 | 0.05 | 0.05 | 0.42 | 0.07 | 16 | 0.57 | 0.06 | 0.06 | 0.04 | 0.37 | 0.06 |
| Slipper limpet | 0.54 | 0.03 | 0.25 | 0.12 | 0.21 | 0.11 | 9 | 0.56 | 0.04 | 0.19 | 0.06 | 0.26 | 0.06 |
| Herring | 0.50 | 0.04 | 0.44 | 0.07 | 0.06 | 0.05 | 6 | 0.59 | 0.04 | 0.24 | 0.07 | 0.17 | 0.06 |
| Acorn barnacle | 0.53 | 0.06 | 0.23 | 0.12 | 0.24 | 0.11 | 2 | 0.43 | 0.04 | 0.28 | 0.08 | 0.30 | 0.07 |
| European smelt | 0.38 | 0.13 | 0.35 | 0.16 | 0.26 | 0.13 | 20 | 0.39 | 0.05 | 0.15 | 0.06 | 0.46 | 0.06 |
| Sole | 0.40 | 0.05 | 0.07 | 0.06 | 0.53 | 0.07 | 24 | 0.39 | 0.04 | 0.05 | 0.03 | 0.56 | 0.05 |
| Baltic clam | 0.40 | 0.04 | 0.16 | 0.11 | 0.44 | 0.10 | 18 | 0.41 | 0.03 | 0.13 | 0.05 | 0.45 | 0.04 |
| Ragworm | 0.35 | 0.05 | 0.55 | 0.09 | 0.09 | 0.07 | 13 | 0.31 | 0.04 | 0.47 | 0.05 | 0.22 | 0.05 |
| Lugworm | 0.35 | 0.07 | 0.08 | 0.07 | 0.57 | 0.08 | 1 | 0.26 | 0.04 | 0.08 | 0.04 | 0.67 | 0.05 |
| Green crab | 0.24 | 0.03 | 0.52 | 0.15 | 0.24 | 0.13 | 3 | 0.31 | 0.03 | 0.29 | 0.05 | 0.40 | 0.04 |
| Cone mud snail | 0.22 | 0.10 | 0.62 | 0.14 | 0.16 | 0.11 | 14 | 0.34 | 0.07 | 0.47 | 0.09 | 0.19 | 0.07 |
| European plaice | 0.17 | 0.05 | 0.75 | 0.09 | 0.08 | 0.06 | 22 | 0.30 | 0.05 | 0.46 | 0.07 | 0.25 | 0.06 |
| European bass | 0.17 | 0.07 | 0.69 | 0.12 | 0.13 | 0.09 | 10 | 0.33 | 0.05 | 0.34 | 0.07 | 0.33 | 0.06 |
| Periwinkle | 0.14 | 0.06 | 0.34 | 0.16 | 0.51 | 0.14 | 17 | 0.16 | 0.04 | 0.32 | 0.05 | 0.53 | 0.05 |
| Sand goby | 0.09 | 0.06 | 0.65 | 0.15 | 0.25 | 0.13 | 12 | 0.20 | 0.05 | 0.41 | 0.07 | 0.39 | 0.07 |
| Shrimp | 0.11 | 0.05 | 0.46 | 0.17 | 0.43 | 0.15 | 7 | 0.20 | 0.04 | 0.24 | 0.05 | 0.56 | 0.05 |
| Eelpout | 0.12 | 0.04 | 0.30 | 0.15 | 0.57 | 0.13 | 25 | 0.26 | 0.05 | 0.15 | 0.06 | 0.59 | 0.06 |
| European flounder | 0.09 | 0.05 | 0.39 | 0.16 | 0.52 | 0.15 | 21 | 0.18 | 0.05 | 0.23 | 0.07 | 0.59 | 0.07 |

Supplementary Figure 1: Map of the Dutch Wadden Sea sample sites including consumers and primary producers sampled for $\delta^{15}N$ of bulk material and amino acids in this study.

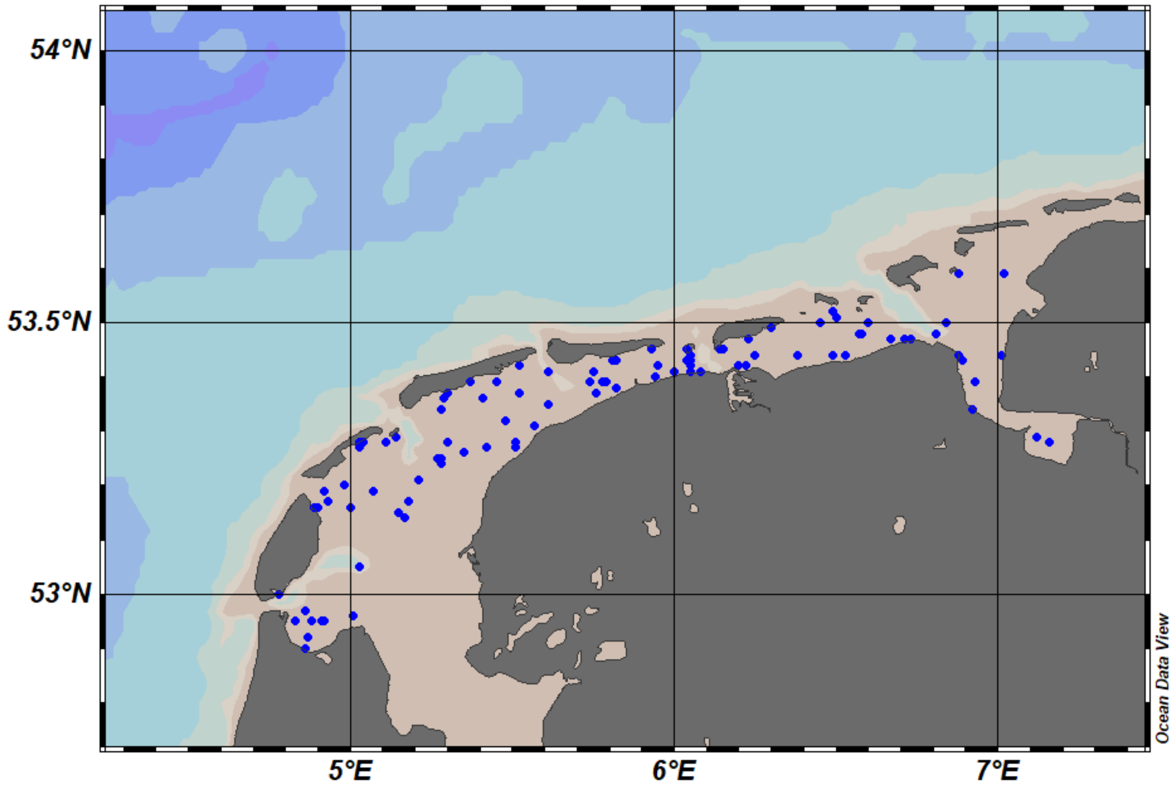

Supplementary Figure 2: Pairs plots for A) two and B) three source mixing models using individuals

**A)**

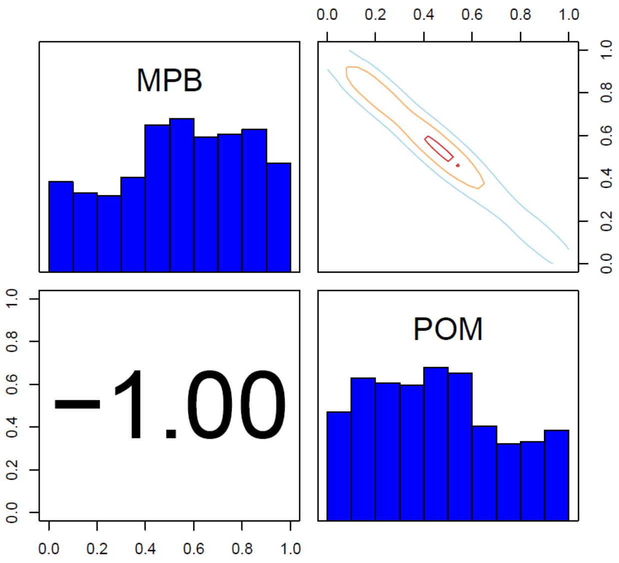


**B)**

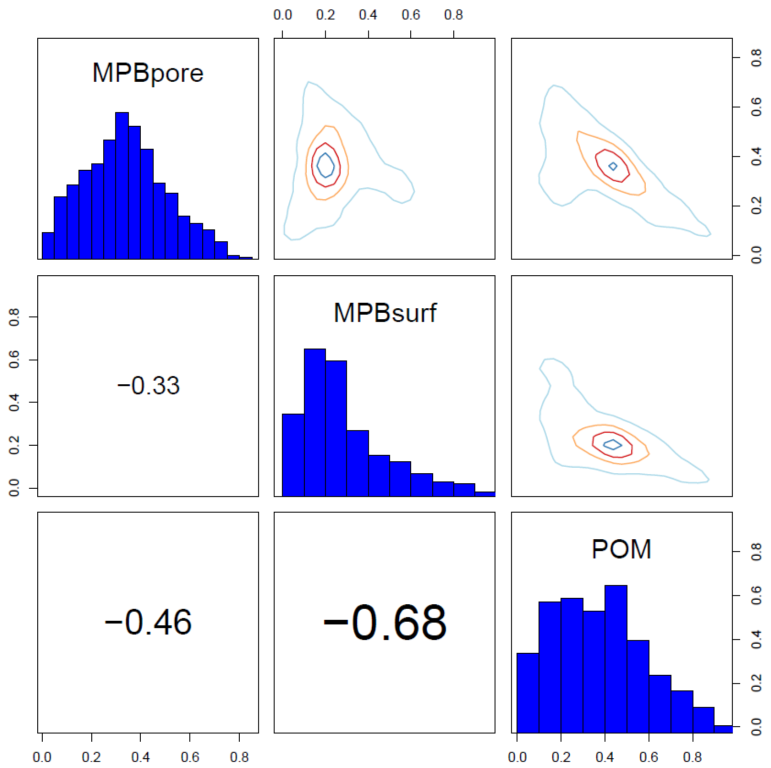
